# A controlled environment assay for the rapid evaluation of verticillium stripe resistance in canola

**DOI:** 10.1101/2025.08.11.669230

**Authors:** Xuehua Zhang, Dale Burns, Kari-Lynne McGowan, Elizabeth Simpson, Jed Christianson, Peter Asiimwe, Pierri Spolti

## Abstract

Verticillium stripe disease is an emerging threat to canola production in Western Canada. Accurately assessing verticillium stripe resistance in breeding germplasm collections is crucial for identifying sources of genetic resistance. Currently, field phenotyping is the most widely used method for evaluating verticillium stripe resistance; however, achieving uniform disease pressure under field conditions presents significant challenges. Here we report a novel controlled environment (CE) soil-less assay for the rapid evaluation of verticillium stripe resistance in canola. The CE results were validated in a field trial which showed a strong correlation between the field data and the CE results. This cost-effective and time-efficient assay enables accurate assessment of verticillium stripe symptoms in a one-month testing cycle.

## Introduction

*Verticillium longisporum* is a soil-borne fungal pathogen that commonly causes wilting, chlorosis, stunting, vascular discoloration, and premature senescence in many plant species, including Brassica species, oat, pea and wheat (Johansson et. al., 2006). Initially, the disease caused by *V. longisporum* in *Brassica napus* was referred to as verticillium wilt. However, since wilting symptoms were not observed in infested oilseed rape or canola plants, the term ‘wilt’ is not applicable. Instead of wilting, stem striping at the beginning of the ripening stage is the typical symptom observed in infested oilseed rape or canola plants. Consequently, verticillium wilt was renamed verticillium stripe in 2016 (Depotter et al., 2016). The disease has become widespread in oilseed rape growing regions in Europe since the 1980s and in canola growing regions in Canada since 2014. Recently, verticillium striping on rapeseed was first reported in northwestern China (Si et al., 2024). Once introduced, the microsclerotia produced by the fungus can survive for 10 to 15 years in soil, even in the absence of host plants. Molecular evidence suggests that *V. longisporum* is a diploid hybrid (allodiploid) and that three species, including *Verticillium dahliae*, have contributed to the genetics of the pathogen. The presence of *V. longisporum* in canola in Manitoba was confirmed by the Canadian Food Inspection Agency (CFIA) in 2014. In 2015, the CFIA reported that *V. longisporum* was most prevalent in canola fields in Manitoba, with lower prevalence observed in Alberta and Saskatchewan fields. Plant disease surveys conducted in recent years indicated outbreaks of verticillium stripe in Manitoba in 2022 and 2024, suggesting that this disease poses a growing threat to canola production (Canadian plant disease survey, https://phytopath.ca/publication/cpds/). The disease can cause significant yield loss if an infection occurs early in the growing season. Estimated yield losses ranging between 10% and 50% under severe disease infection were reported (Dunker et al. 2008), one of the most critical factors affecting yield loss being the reduction in seed size and lipid content. Verticillium stripe often coexists with other diseases, such as blackleg and sclerotinia stem rot. When plants were affected by both *Leptosphaeria maculans*, the causal agent of blackleg, and *V. longisporum*, higher yield losses were observed (Wang et al., 2023).

Verticillium stripe is a monocyclic disease, i.e. it has only one disease cycle each year. The fungus can enter the vascular system through the roots via fungal hyphae. Subsequently, conidia are produced locally in the xylem and transported through the plant’s vascular system, disrupting the normal flow and functionality of water and nutrient transport. This ultimately leads to the plugging, blackening, collapse, and shriveling of the xylem. Disease symptoms in canola include leaf chlorosis, early ripening, and stunting. As the disease progresses, symptoms can include necrosis and shredding of stem tissue. Once the plant is fully ripe, the stem may peel away to reveal tiny black microsclerotia that resemble ground pepper.

Currently, crop rotation with non-host crops and effective weed management are the most common practices for managing this disease. Although the transmission of this pathogen occurs through microsclerotia present in the soil and plant debris from the previous growing season, Zhang et al. (2019) reported the potential for seed transmission of this pathogen in *B. napus*. While genetic resistance may be one of the most effective methods of disease control, the understanding of genetic resistance in commercial canola hybrids is still very limited. Although Wang et al. (2024) reported the identification of quantitative trait loci (QTLs) contributing to verticillium stripe resistance, it takes a considerable amount of time to introduce the genetic resistance into breeding lines. Although seed developers and public researchers have begun working on the identification of genetic resistance in Brassica species in recent years, it still takes a significant amount of time for seed developers to identify sources of genetic resistance and to breed for genetic resistance. Phenotyping is crucial for helping breeding programs accelerate the breeding process.

Field phenotyping is the most common method for screening plant material for disease resistance; however, the data return rate from field trials can be unpredictable due to environmental conditions, inoculation techniques, and agronomic practices. Additionally, scaling up field trials is challenging, with only one growing season per year to conduct these trials. Researchers have developed several CE screening protocols and inoculation techniques for verticillium stripe, demonstrating that CE disease screening is more repeatable than field trials (Eynck et al., 2008; Cui et al., 2023). However, these methods tend to be labor-intensive and resource-demanding, making it difficult to apply them in large-scale breeding germplasm screening. Therefore, the development of an efficient and predictive CE germplasm screening protocol is critical for disease resistance breeding. The objectives of this study were to develop an efficient CE assay to screen verticillium stripe resistance and to understand the relationship between data generated under CE and field conditions.

## Material and Methods

### Controlled environment (CE) disease screening

#### Plant Material

A total of 162 *Brassica napus* germplasm collection accessions, including the susceptible control variety Westar, were screened under CE conditions. A subset of 28 accessions was screened using 6 replicates in a separate CE experiment and evaluated in field trials to understand the relationship between seedling testing results in the CE and adult plant assessments under field conditions.

#### Material Preparation

The regular-sized CYG germination pouch (Mega International, https://mega-international.com/) was used for CE verticillium stripe screening. Although the recommended lifespan of the germination pouch is two weeks, we successfully grew plants in the pouch for a full month. Before seeding, 50 mL of tap water was added to the bottom of each pouch and the pouch was placed in the polypap stand. The polypap stands were placed in a 10 × 20-inch plastic tray. Approximately 15 seeds were placed in the trough formed by the paper wick. Each entry was seeded in two pouches, with each pouch considered one replicate. The susceptible check variety Westar was repeated six times, and an additional six Westar replicates were seeded as mock controls. Two extra Westar pouches were included in each tray to monitor the progress of disease infection. After seeding, the pouches were kept at 4°C in the dark for 2 days to enhance the rate and synchrony of germination. Subsequently, the pouches were transferred to a growth chamber and grown at 24°C with a 16-hour photoperiod.

#### Inoculum Preparation

One pure isolate (V20RSH2) of *Verticillium longisporum* was used to produce inoculum. This isolate originated from diseased canola plants in a field near Rosenort, Manitoba in 2020, and belongs to the A1/D1 group as determined by the methods described by Zou et al. (2020). The isolate was preserved on paper discs containing mycelium and stored at -20°C.

One paper disc was plated onto a potato dextrose agar (PDA) plate (9-cm diameter petri dish) and incubated in the dark at room temperature for 14 days to one month. Once the mycelium covered over 80% of the plate surface, the entire plate was cut into plugs using a scalpel and placed into a 1 L flask containing 0.5 L potato dextrose broth (PDB) media. The flasks were shaken at room temperature (24°C) at approximately 150 rpm under constant fluorescent lighting for 14-21 days to achieve a spore concentration of greater than 1 × 10^7^ spores/mL. The resulting spore suspension was then filtered using sterile Miracloth. The spore concentration was determined using a hemocytometer under a compound microscope (400× magnification) and diluted to 1 × 10^7^ spores/mL.

#### Plant inoculation

Ten days after seeding, the water in the germination pouch was removed, and the roots were gently wounded using a 1 mm microneedling roller. The roller was positioned along the seeding line and rolled for two complete passes. Approximately 20 mL of liquid inoculum was then added to the pouch, the wounded roots were submerged in the inoculum for 1 hour. For mock-inoculated control, the plants were submerged in sterilized purified water. After inoculation, the inoculum was removed, and 20 mL of tap water was added.

#### Plant maintenance

After inoculation, the plants were grown at room temperature (24°C) under fluorescent lighting with a 16-hour photoperiod. Two days post-inoculation, the water was removed, and 20 mL of liquid ½ MS medium (Murashige & Skoog Basal Medium with Vitamins, PhytoTech Labs) was added to each pouch. One week later, an additional 20 mL of liquid ½ MS medium was added to each pouch. The liquid at the bottom of the pouch was monitored periodically and refilled with tap water as needed. Two weeks after inoculation, the pouches were misted with tap water daily to prevent drying out. Disease symptoms were monitored starting two weeks post-inoculation, and disease severity was evaluated when severe symptoms appeared on the susceptible check, typically occurring 21 days post-inoculation. The procedure of the CE assay is demonstrated in Figure 1.

**Figure 1.**
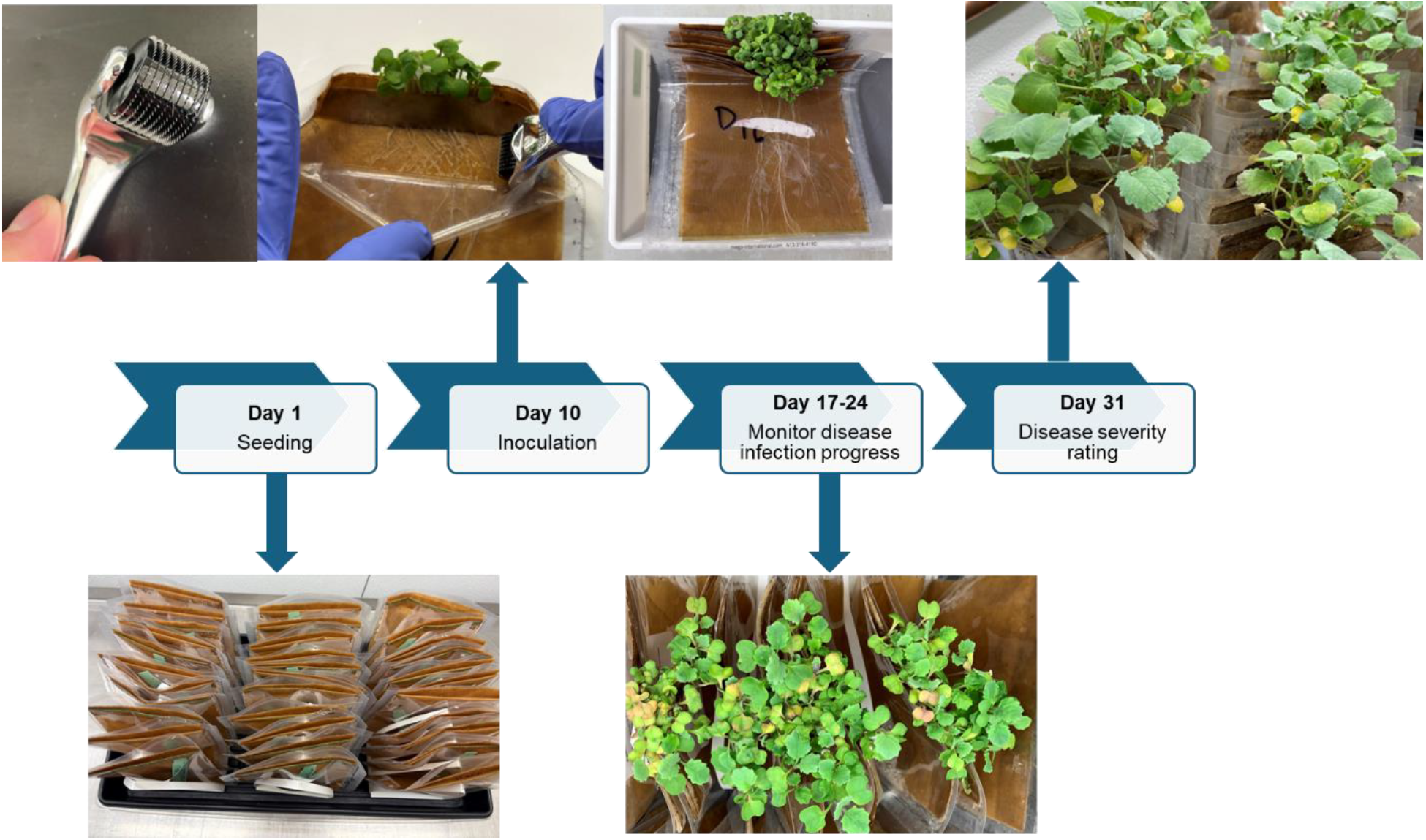
Workflow of the controlled environment (CE) assay for evaluating verticillium stripe resistance in canola.

### Disease severity evaluation and data collection

Plants were assessed for disease severity at 21 days post-inoculation using a modified 1-9 disease severity rating scale based on Zeise (1992) and Eynck et al. (2009) (Figure 2.). Mock-inoculated controls were used as comparators to evaluate the degree of stunting and chlorosis in the check plants. The mean disease severity score for each entry was calculated, and entries were categorized as follows: Resistant (R) if the mean score was < 4.5; Moderately Resistant (MR) if 4.5 < mean < 5.5; Moderately Susceptible (MS) if 5.5 < mean < 6.5; and Susceptible (S) if the mean score was > 6.5.

**Figure 2.**
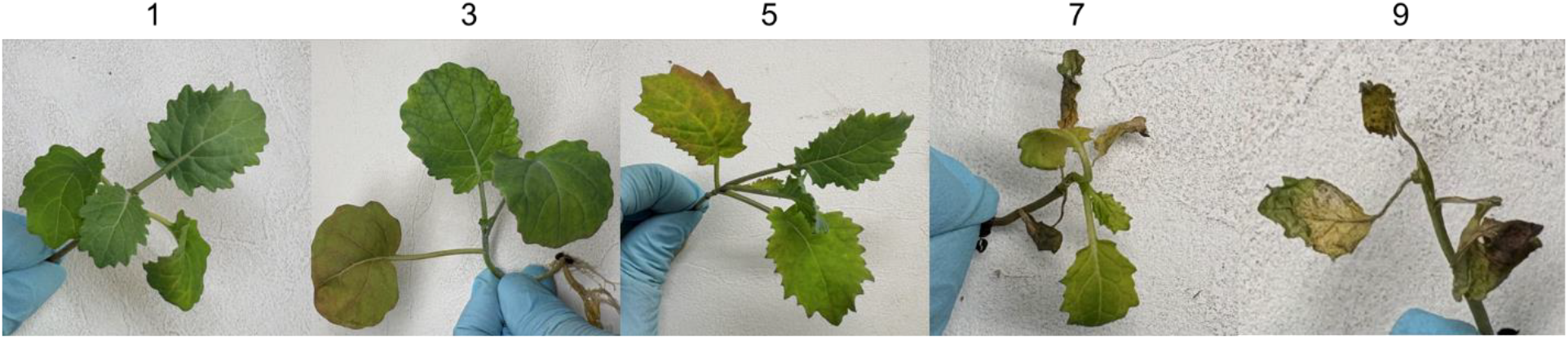
Disease rating scale for assessing symptoms caused by *Verticillium longisporum* at the seeding stage. 1 = No visible symptoms; 3 = Slight symptoms on older leaves; 5 = More than 50% of the leaves showing symptoms; 7 = More than 75% of the leaves showing symptoms; 9 = The plant is dead.

### Field trial

To validate the CE disease screening protocol, a subset of 28 *B. napus* accessions representing all four resistance categories was tested under field conditions.

The field trial was conducted in 2024 at two sites in Manitoba that were previously infested with Verticillium stripe. The plot size for the field trial at the first site was 1 meter by 1.5 meters and 1.5 meters by 5 meters at the second site. All trials were planted using a yield trial planter and maintained according to standard agronomic practices for canola production in Canada. The trials at both locations utilized a randomized complete block (RCB) design. Each entry was replicated three times in both trials.

Disease severity was assessed after harvest, based on both plot level and individual plant level infection (Figure 3.). The assessment utilized a 1-9 rating scale modified from Dunker et al. (2008) and Steventon et al. (2002). For the individual plant level assessment, approximately 25 stubbles were randomly pulled from each plot, and the average disease severity rating was calculated. To align with CE disease screening results, the mean disease severity score for each entry was calculated, and entries were categorized as follows: Resistant (R) if the mean score was < 4.5; Moderately Resistant (MR) if 4.5 < mean < 5.5; Moderately Susceptible (MS) if 5.5 < mean < 6.5; and Susceptible (S) if the mean score was > 6.5.

**Figure 3.**
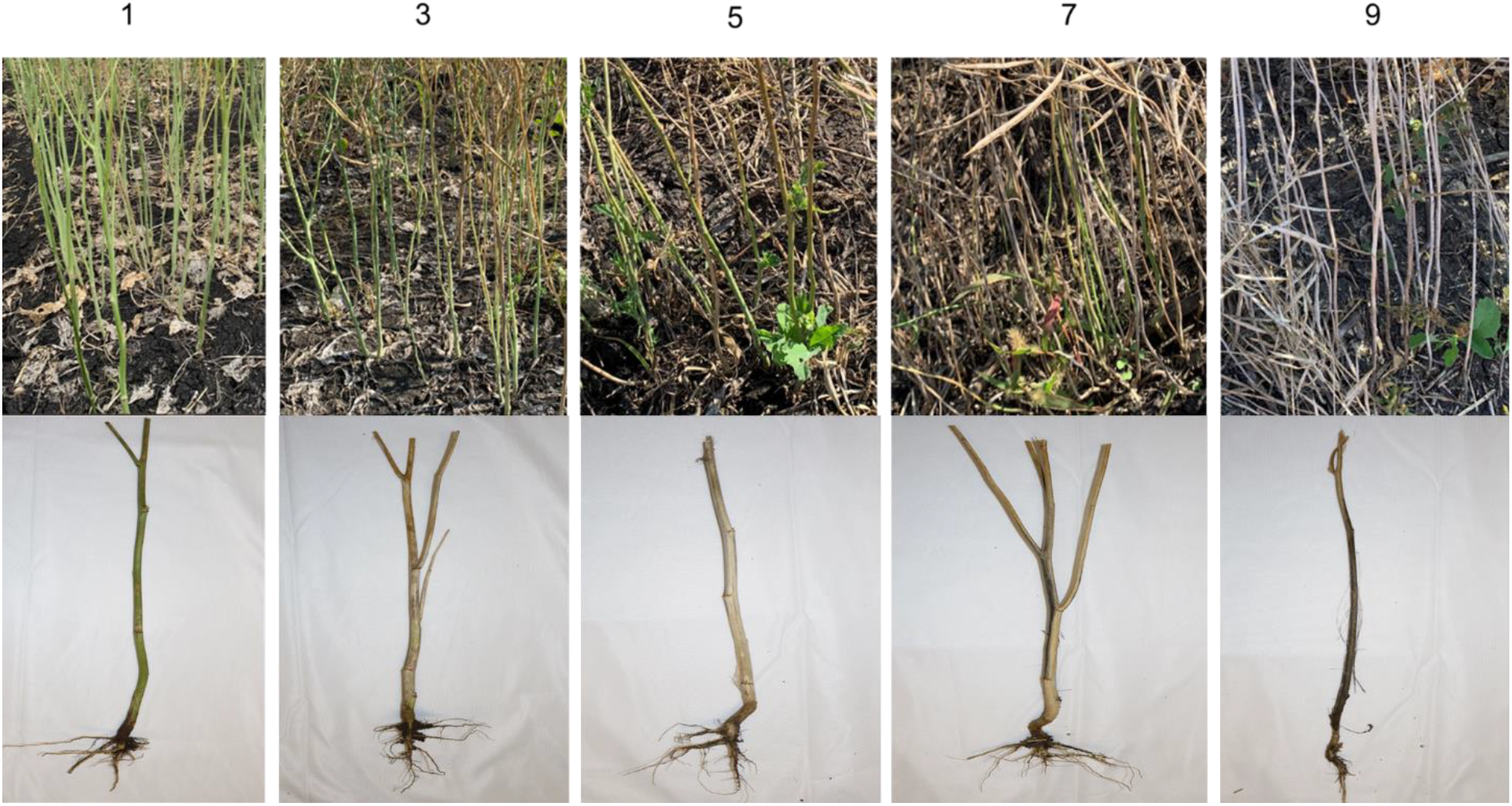
Verticillium stripe disease severity rating scale for field trials. For individual plots: 1 = No infestation; 3 = Low infestation: <30% of plants showing symptoms; 5 = Medium infestation: 30-50% of plants showing symptoms; 7 = Strong infestation: 50-70% of plants showing symptoms; 9 = >70% of plants showing symptoms. For Individual Plants: 1 = No infestation; 3 = Low infestation: Streaky discoloration of the stem; sporadic striping of the epidermis; microsclerotia in up to 25% of the epidermis; stem pith or single branches completely colonized with microsclerotia; 5 = Medium infestation: Epidermis easy to detach from the stem; up to 50% of the stem or stem pith colonized with microsclerotia; 7 = High infestation: Complete discoloration of the stem; epidermis fibrous or only fibrous remnants of the epidermis present; up to 75% of the stem colonized with microsclerotia; 9 = Severe infestation: Premature dying off; complete discoloration of the stem interspersed with microsclerotia (>75%).

## Results

### Controlled environment (CE) Verticillium stripe phenotyping

All 162 entries were tested in a single screen run. Disease symptoms began to appear approximately 7 to 10 days after inoculation, with significant disease progression observed at around 18 to 21 days post-inoculation. The disease severity rating was conducted on 21 days post inoculation. For each replicate, 10-15 individual plants were assessed for disease severity and the average disease severity rating score was calculated. The mean disease severity was calculated per entry. Overall, a variable level of verticillium stripe resistance was observed among the different entries and the variation between the two replicates was minimal, with mean disease severity ratings ranging from 3.5 to 8.6 and standard deviations ranging from 0 to 0.5. Data collected from the two replicates was highly correlated, suggesting a high level of repeatability (Figure 4.). The susceptible check variety Westar was confirmed as one of the most susceptible accessions. Out of the 162 entries evaluated in the study, 16 exhibited resistance to *V. longisporum*, 47 were moderately resistant, 68 were moderately susceptible, and the remaining 31 entries were highly susceptible to the pathogen (Figure 5.). Symptoms exhibited by entries in different resistance categories were visually distinguishable (Figure 6.).

**Figure 4.**
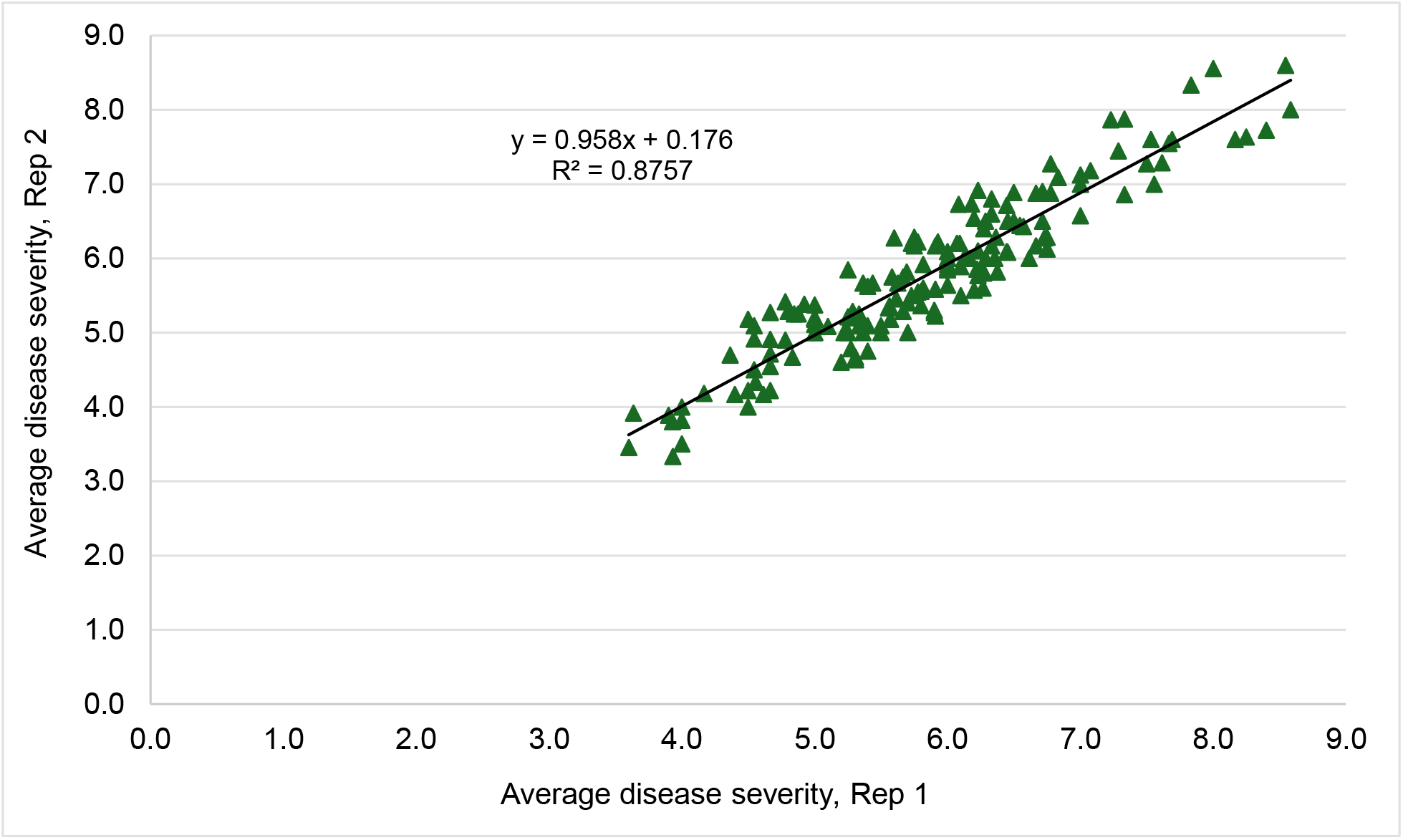
The correlation between two replicates of controlled environment screening.

**Figure 5.**
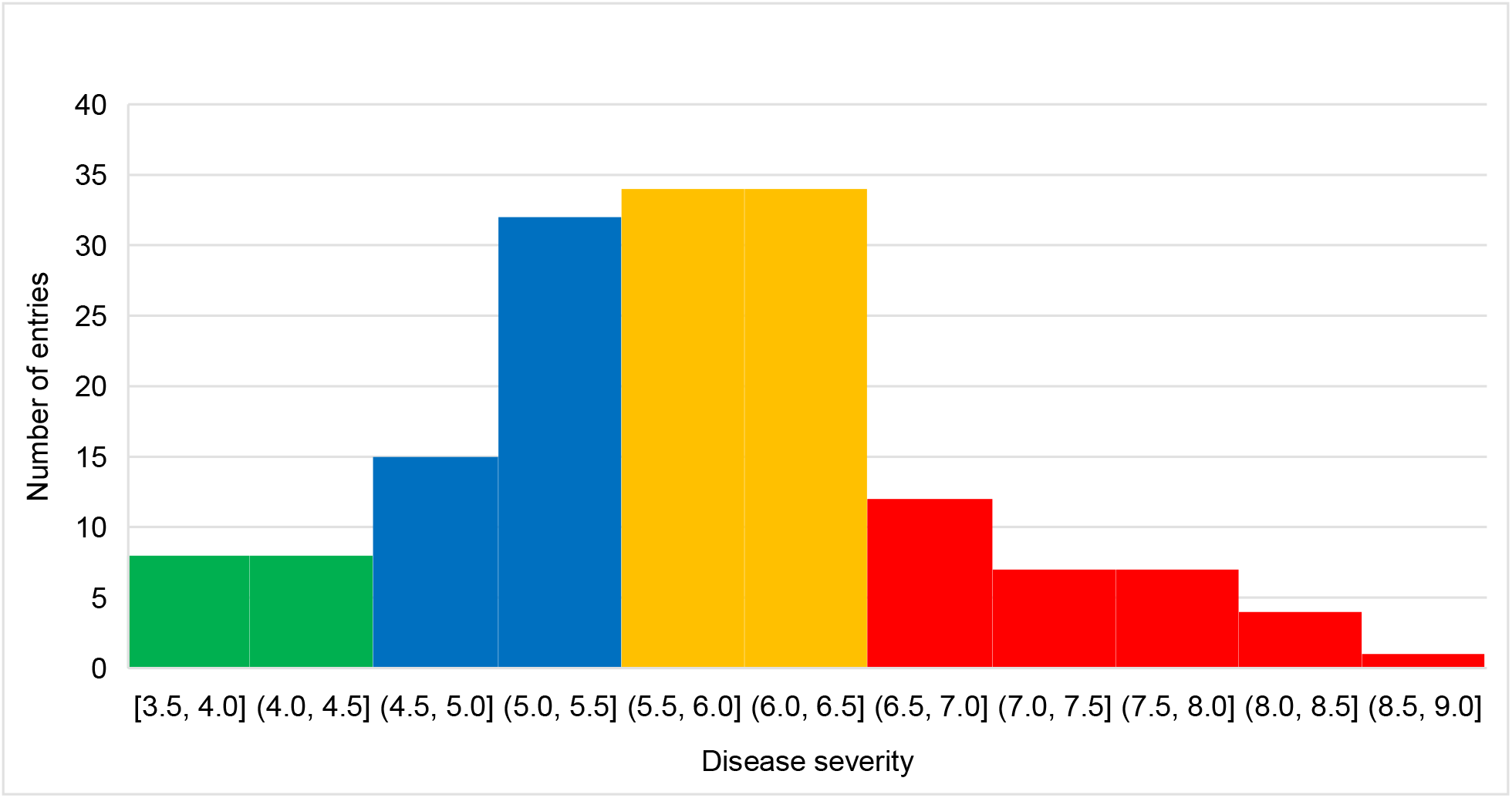
Distribution of entries across different disease severity categories. Green bars representing resistant (R) entries, blue bars representing moderately resistant (MR) entries, orange bars representing moderately susceptible (MS) entries, and red bars representing susceptible (S) entries.

**Figure 6.**
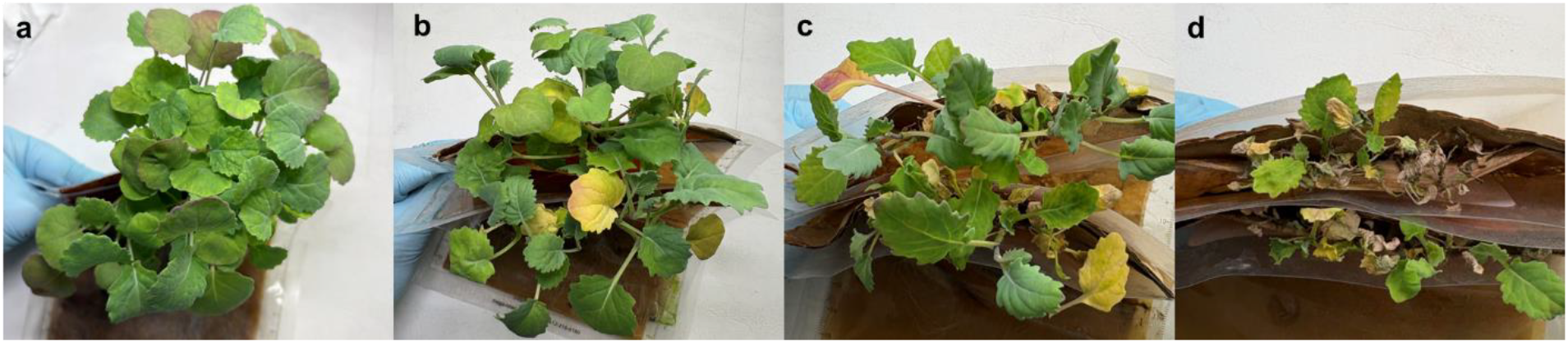
Symptoms on entries are categorized as resistant (a), moderately resistant (b), moderately susceptible (c) and susceptible (d) canola entries.

A subset of 28 entries, including the susceptible check, was screened in another run with 6 replicates to observe variations and verify results from the first screening run. The second run confirmed the findings from the first run, and the results were highly correlated (data not shown).

### Field verticillium stripe phenotyping

Disease pressure was very low at the first site due to early-season flooding, while the disease pressure at the second site was sufficient to differentiate between entries. Consequently, data collected from the first site is not discussed in this paper. Data from the second site was included in the analyses. Overall, disease pressure at the Oakville site was evenly distributed across the field. The average disease severity data collected at the plant level were more variable than the data collected at the plot level; therefore, the plot-level disease severity data were used for further analysis.

### Comparison between controlled environment (CE) verticillium stripe screening data and field verticillium stripe testing data

The CE disease severity ratings ranged from 4.19 to 7.96, with standard deviations ranging from 0.08 to 0.84. In contrast, field disease severity ratings ranged from 5.00 to 9.00, with standard deviations from 0.00 to 2.65. The variation between replicates was significantly higher in the field trial, primarily due to environmental factors and suspected uneven distribution of inoculum in the field. The CE disease severity data and field data were positively correlated (Figure 7), suggesting that CE data can be used to predict field disease risk.

**Figure 7.**
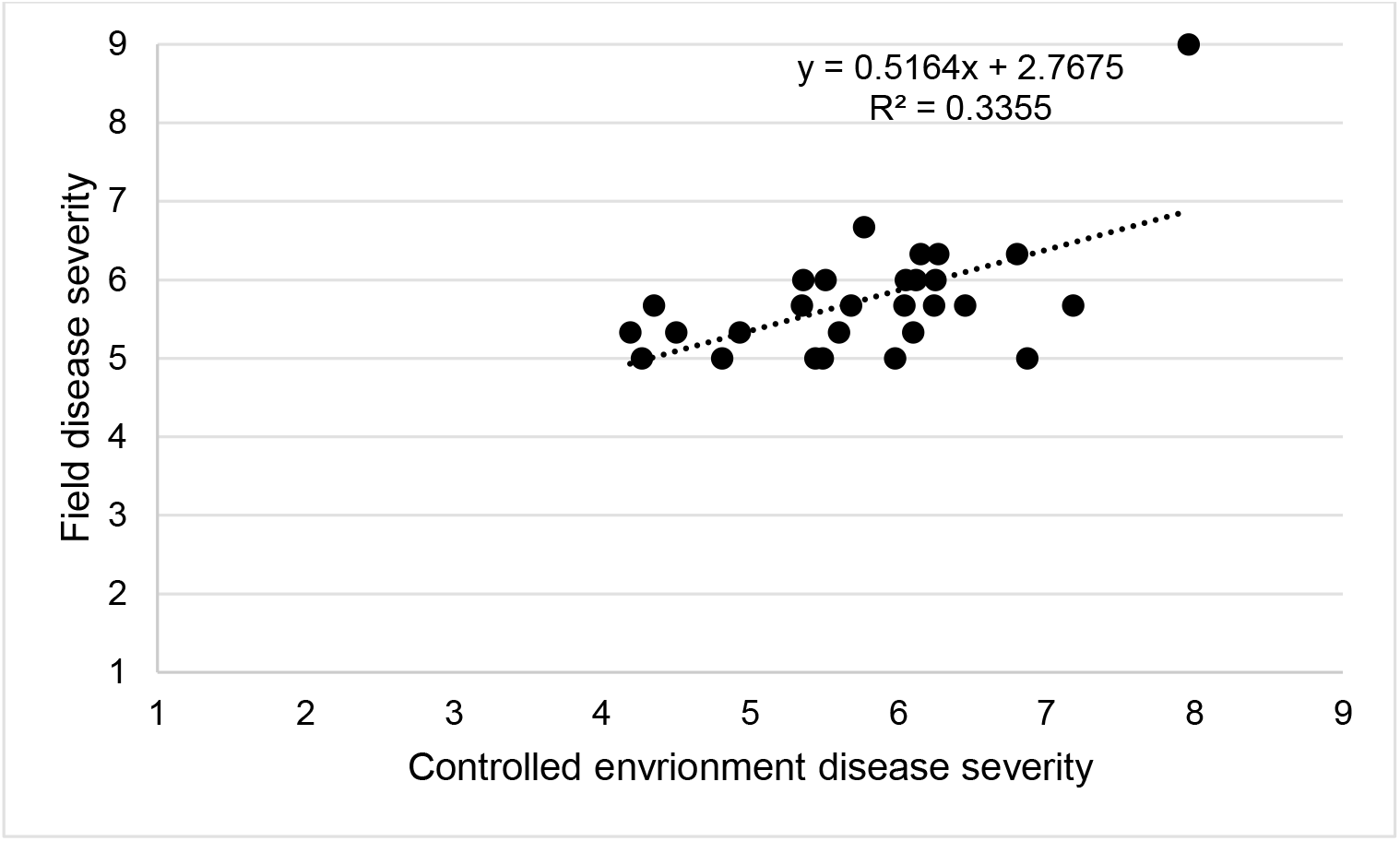
The correlation between controlled environment (CE) disease severity and field disease severity.

Out of the 28 entries tested in the field environment, the results for 17 entries aligned well with the CE testing results, with these entries categorized into the same disease resistance category using both data sources. The most susceptible entry in the CE testing was also the most susceptible entry in the field environment. Five entries were found to be more susceptible in the field trial. Specifically, Bayer 07 and Bayer 26 were classified as R in the CE but were categorized as MR in the field trial. Bayer 27 was classified as R in CE but was MS in the field trial. Bayer 02 was categorized as MR in the CE, while the field trial suggested it was MS. Bayer 11 was classified as MS in the CE but was found to be susceptible in the field trial. Conversely, six entries showed greater resistance in the field trial. Bayer 20, Bayer 21, and Bayer 23 were categorized as MS in the CE but as MR in the field trial. Bayer 04 and Bayer 10 were classified as S in the CE but as MS in the field trial. Bayer 08 was classified as S in the CE but as MR in the field trial. The comparison between the CE data and the field-testing data was summarized in Table 1.

**Table 1.** Summary of controlled environment (CE) and field Verticillium stripe disease severity ratings and resistance categories: Data highlighted in light green represents entries categorized into the same disease resistance category. Data highlighted in light blue indicates entries that were more susceptible in the field environment. Data highlighted in light grey represents entries that were more resistant in the field environment.

|  | Controlled environment |  | Field environment |  |
| --- | --- | --- | --- | --- |
| Accessions | Mean disease severity and standard deviation | Resistance category | Mean disease severity and standard deviation | Resistance category |
| Bayer01 | 7.96 ± 0.37 | S | 9.00 ± 0.00 | S |
| Bayer02 | 5.35 ± 0.76 | MR | 5.67 ± 0.58 | MS |
| Bayer03 | 4.93 ± 0.49 | MR | 5.33 ± 0.58 | MR |
| Bayer04 | 7.18 ± 0.84 | S | 5.67 ± 1.15 | MS |
| Bayer05 | 6.45 ± 0.34 | MS | 5.67 ± 0.58 | MS |
| Bayer06 | 6.05 ± 0.84 | MS | 6.00 ± 1.00 | MS |
| Bayer07 | 4.19 ± 0.13 | R | 5.33 ± 0.58 | MR |
| Bayer08 | 6.87 ± 0.53 | S | 5.00 ± 2.65 | MR |
| Bayer09 | 6.27 ± 0.64 | MS | 6.33 ± 1.15 | MS |
| Bayer10 | 6.80 ± 0.77 | S | 6.33 ± 1.53 | MS |
| Bayer11 | 5.77 ± 0.68 | MS | 6.67 ± 0.58 | S |
| Bayer12 | 5.44 ± 0.25 | MR | 5.00 ± 1.00 | MR |
| Bayer13 | 4.50 ± 0.59 | MR | 5.33 ± 1.53 | MR |
| Bayer14 | 6.15 ± 0.56 | MS | 6.33 ± 0.58 | MS |
| Bayer15 | 6.04 ± 0.23 | MS | 5.67 ± 2.08 | MS |
| Bayer16 | 6.25 ± 0.51 | MS | 6.00 ± 0.00 | MS |
| Bayer17 | 5.68 ± 0.29 | MS | 5.67 ± 1.53 | MS |
| Bayer18 | 5.49 ± 0.28 | MR | 5.00 ± 0.00 | MR |
| Bayer19 | 5.51 ± 0.58 | MS | 6.00 ± 0.00 | MS |
| Bayer20 | 5.60 ± 0.08 | MS | 5.33 ± 0.58 | MR |
| Bayer21 | 6.10 ± 0.65 | MS | 5.33 ± 1.15 | MR |
| Bayer22 | 6.12 ± 0.31 | MS | 6.00 ± 1.73 | MS |
| Bayer23 | 5.98 ± 0.52 | MS | 5.00 ± 1.00 | MR |
| Bayer24 | 6.24 ± 0.42 | MS | 5.67 ± 0.58 | MS |
| Bayer25 | 4.81 ± 0.08 | MR | 5.00 ± 1.00 | MR |
| Bayer26 | 4.27 ± 0.21 | R | 5.00 ± 1.00 | MR |
| Bayer27 | 4.35 ± 0.36 | R | 5.67 ± 0.58 | MS |
| Bayer28 | 5.36 ± 0.78 | MR | 6.00 ± 1.00 | MS |

## Discussion

The continued spread of verticillium stripe in Canada necessitates breeding efforts to eliminate highly susceptible hybrids at early stages of development and to develop commercial hybrids that exhibit resistance or moderate resistance to this pathogen. The availability of efficient and effective phenotyping protocols is crucial for characterizing breeding materials and for developing breeding tools, such as genomic prediction models and molecular markers, to assist in breeding selection processes. Previous CE screening methods for verticillium stripe reported in the literature have been resource-intensive, time-consuming, and labor-intensive (Eynck et al., 2009; Cui et al., 2023). For example, Eynck et al. (2009) screened 1348 Brassica accessions in 26 independent screening runs over more than two years, and 20 inoculated seedlings per accession were assessed. In contrast, this study screened 162 entries, evaluating approximately 25 to 30 individual seedlings per entry in two replicates within a one-month timeframe. The root-dip inoculation method has been widely employed in various CE screening methods for verticillium stripe. Moreover, wounding the roots is essential to facilitate the penetration of the pathogen. Traditional methods of root wounding, such as root tip excision, can introduce variability between plants and replicates. To address this issue, we improved the root-dipping technique by employing microneedling rollers to create uniform wounding sites on the roots. The CE screening assay described in this study integrates procedures and techniques that have proven effective in other protocols while adopting a soilless testing approach to optimize space and resource utilization. This assay enables the screening of many entries in a single testing cycle, utilizing minimal growth area, and demonstrates consistent data across replicates and independent experiments.

The screening of germplasm collections in this study provides valuable data for further exploration. While some entries were highly susceptible to the pathogen, a variable level of resistance was observed in a significant number of entries. Strong resistance to *V. longisporum* was identified in several accessions, which may serve as potential donor lines for future breeding projects. This CE assay enabled us to screen large populations using minimal seed quantities and growing resources within a one-month testing cycle.

Field trial was conducted to assess whether the data obtained from CE screenings can be used to predict field performance. Compared to field testing data, CE testing data is more repeatable, with standard deviations significantly lower than those of the field-testing results. Although some studies suggest that results from CEs do not easily translate to field responses (Eynck et al. 2009), this study demonstrated a reasonable correlation between CE data and field-testing data. Specifically, most entries categorized as susceptible (S) or moderately susceptible (MS) in the CE were also classified as S or MS in the field environment. It is possible that disease pressure in the field is sufficiently high to differentiate the more susceptible entries. However, controlling the factors affecting disease infection in the field is challenging, and the distribution of inoculum is often uneven. Several isolates were purified from infested stubbles collected in the second testing site, and pathogenicity testing was conducted to compare the aggressiveness of these isolates with the isolate used in CE testing. The results indicated that these isolates were more aggressive than V20RSH2. This finding helps explain why some entries were classified as resistant (R) in CE testing but as moderately resistant (MR) or moderately susceptible (MS) under field conditions. Additionally, variations in the aggressiveness of isolates originating from different fields could be another factor influencing the correlation between CE data and field data.

For field disease severity ratings, we found that plot-level ratings based on the percentage of infested plants were more consistent between replicates and more efficient. In contrast, plant-level ratings were time-consuming and labor-intensive, with greater variability in the data due to sampling issues. It was particularly challenging to extract heavily infested plants, as they are fragile, leading to potential subsampling bias.

Currently, there are several different verticillium stripe disease severity rating scales used by various research groups (Eynck et al., 2009; Cui et al., 2023; Wang et al., 2023). While each rating scale has its own advantages, comparing disease severity data across different organizations is challenging, making it difficult for growers and agronomists to understand the relationship between the data and disease risk. In this study, we found that utilizing a 1-9 scale for both CE disease severity evaluation and field disease severity assessment is effective. A 1-9 scale has been widely used for many other plant diseases, facilitating easier data analysis and comparison of disease severity across different diseases. The development of an industry-standard verticillium stripe disease severity rating scale in Western Canada is essential.

## Conclusions

The CE assay reported in this paper is highly reproducible and saves both time and resources. While this CE verticillium stripe screening assay can accurately assess resistance and susceptibility to *V. longisporum*, field validation remains essential to confirm these resistance levels. A combination of results from CE testing and field testing provides a highly reliable assessment of verticillium stripe resistance. Using CE testing data to predict field disease risk is reliable and can be implemented early in the breeding pipeline. Although entries categorized as R or MR in CE may not necessarily exhibit the same resistance under field conditions due to the aggressiveness of the isolates present in the field and various environmental factors, MS and S entries identified using this assay tend to be more susceptible in field conditions and carry a higher disease risk. This assay can be employed to eliminate highly susceptible entries at an early breeding stage, thereby mitigating the risk of high susceptibility to this disease in field conditions.

## Acknowledgements

We would like to acknowledge Reginald Morton, Shuying Xiong, and Teresa Reyes for their valuable technical assistance in CE assays. We also extend our gratitude to Adam Meisner and Paul Gregoire for their technical assistance in field trials.

